# Pre-Germinal Center Interactions with T Cells are Natural Checkpoints to Limit Autoimmune B Cell Responses

**DOI:** 10.1101/2022.07.20.500862

**Authors:** Kate A Parham, Xiu Xia Sherry Tan, Daniel M Morelli, Lika Chowdhury, Heather C Craig, Steven M Kerfoot

## Abstract

Interactions with antigen-specific T cells drive B cells activation and fate choices that ultimately determine the quality of high-affinity antibody responses. As such, thse interactions, and especially the long-lived interactions that occur prior to germinal center formation, may be important checkpoints to regulate undesirable responses. We directly observed interactions between T and B cells responding to the self-antigen Myelin Oligodendrocyte Glycoprotein (MOG) and found that they are of lower quality compared to interactions between cells responding to the model foreign antigen NP-ovalbumin (NP-OVA). This was associated with reduced expression of molecules that facilitate these interactions on the B cells but not on T cells. B cell expression of these molecules was not dictated by the T cell partner, nor could the relative lack of expression on MOG-sp. B cells be reversed by a multivalent antigen. Instead, MOG-sp. B cells were inherently less responsive to B cell Receptor stimulation than MOG-non-sp. cells. However, the phenotype of MOG-sp. B cells was not consistent with previous descriptions of autoimmune B cells that had been tolerized via regular exposure to systemically-expressed self-antigen. This suggests that alternate anergy pathways may exist to limit B cell responses to tissue-restricted self-antigens.

## Introduction

Interactions between antigen-specific (Ag-sp.) T cells and B cells are the foundation of the Germinal Centre (GC) response, as T cells provide essential survival and differentiation signals to B cell Receptor (BCR)-stimulated B cell clones (1–3). Because of their critical nature, these interactions represent potential opportunities to modulate the quality of the humoral response to a given Ag. Similarly, these interactions also represent a potential checkpoint to regulate and limit undesirable humoral responses to self-Ag.

Model Ag Systems are experimental toolkits that combine a manipulatable Ag with Ag-sp. T and B cell clones that can be identified and tracked throughout the immune response. We and others have used Model Ag Systems to characterize the timeline and kinetics of the interactions between T cells and B cells as they occur during the development of the GC (reviewed in (4)). Naïve B cells first engage Ag with their BCR while in the follicle, but their full activation depends on additional signals from activated T cells (reviewed in (1)). To access these signals, Ag-stimulated B cells relocate to the follicular border where they encounter pre-T Follicular Helper (Tfh) clones that themselves were recently activated through interaction with mature Ag-presenting Dendritic Cells in the T cell Zone. The B cells present Ag that they internalized via their BCR to these pre-Tfh cells, resulting in long term (up to hours-long) cognate interactions (5, 6) through which the B cells become activated to proliferate. Therefore, these initial interactions, which occur between 1 and 3 d post exposure to Ag, represent a critical checkpoint that ensures that Ag-stimulated B cells only become fully activated in the presence of a cognate T cell response.

In addition to this gate-keeping role, pre-GC interactions between cognate T cells and B cells are central to determining the quality of the resulting humoral response (reviewed in (1, 7)). In addition to inducing B cell proliferation, they also drive early fate decisions of activated B cell clones to differentiate to either pre-GC Memory B cells, short-lived IgM-producing Plasmablasts, or to re-enter the follicle to initiate GC formation. More recently it was recognized that these early pre-GC interactions also drive isotype class switch (8), which is a fundamental step in the optimisation of the humoral response to a given antigen. Once the GC has formed, B cell survival and maintenance of the GC, as well as differentiation into Memory or Plasma Cell subsets, remains dependent on interactions with cognate Tfh cells (reviewed in (2)), but these interactions are typically much shorter in duration than those that occur pre-GC at the follicular border.

We previously compared GC responses initiated by two very different Model Ags (9); Nitrophenyl -haptenated Ovalbumin (NPOVA), a standard foreign Model Ag; and Myelin Oligodendrocyte Glycoprotein (MOG), a Central Nervous System (CNS)-specific Model Self-Ag. We identified considerable differences in the outcome of the GC response initiated by these Model Ags; particularly differences in the outcome of the early, pre-GC T cell/B cell interactions. Indeed, there was less initial B cell proliferation, virtually no early short-lived Plasmablast differentiation, and much reduced class switch in the MOG response compared to the NPOVA response. While the GCs themselves formed with normal kinetics in response to immunization with MOG, we found that the outcomes of Tfh/B cell interactions in the mature GC were also defective in that they could not sustain the GC. B cell memory, whether generated pre-GC or from the mature GC, was not long-lived. Despite this, anti-MOG antibodies were measurable in the circulation of MOG-immunized mice, including class switched IgG at levels comparable to anti-NP antibodies following NPOVA immunization. Therefore, despite the considerable differences in how the MOG-GC developed and its failure to produce effective memory, the antibody response itself was not entirely defective.

Most investigation of B cell responses to self-Ags have focussed on systemically-expressed Ags, such as anti-DNA Ag systems or transgenic systems where an otherwise-foreign Model Ag is expressed ubiquitously. These studies revealed tolerance mechanisms that limit B cell responses to self-Ags (reviewed in (1, 10–12)), including Central Tolerance deletion of autoreactive cells or BCR editing of receptors that bind self-Ag with high affinity, and also peripheral tolerance of B cells that recognize common systemic antigen that are typically associated with repeated BCR stimulation in the absence of T cell help.

Unlike these Ags, the expression of MOG is restricted to the CNS and not present in lymphoid tissues where B cells develop (13). Nevertheless, it remains a highly relevant B cell autoantigen for investigation. Immunization with MOG-derived peptide is well known to result in T cell-driven autoimmune pathology termed Experimental Autoimmune Encephalomyelitis (EAE), which is often used to model human Multiple Sclerosis (MS) (14). Immunization with a larger conformational MOG protein allows for the possibility of B cells participating in the autoimmune response (15), however the role of anti-MOG B cells in either EAE or MS is not clear. EAE develops spontaneously in mice with enriched T cell and B cell specificity for MOG (16, 17), and therefore MOG-sp. B cells contribute to disease initiation in this model. Anti-MOG B cells are also necessary for disease induced in mice through immunization with human MOG protein Ag (18, 19). In contrast, anti-MOG B cells are not necessary to initiate disease induced by rodent MOG (19), but we have found they can contribute to disease course and pathology (15). The presence of anti-MOG antibodies has recently been determined to be the distinguishing feature of a subset of human patients with demyelinating disease recently recognized as a separate disorder overlapping with MS and related diseases (20). B cells do accumulate in the meninges, but only rarely in the CNS parenchyma, of both human MS patients (21–24) and select mouse EAE models (15, 25–27). However, we and others have shown that the meningeal B cells in these mice are largely not specific for MOG-antigen (28, 29), and a significant proportion of Plasma Cells in the inflamed CNS in both animal models and humans are similarly not specific for autoantigen (30). Therefore, while it is clear from clinical trials that B cell depletion limits disease progression in MS patients (31) and some EAE models (our unpublished observations), the role of autoimmune B cells specifically and the anti-MOG GC in ongoing disease is not clear.

Our previous investigation of the MOG GC (9) suggests that, while the GC response is not absent following immunization, it is attenuated compared to a foreign Model Ag. We found that this could be only partly attributed to the T cell partner in the response, and that there was also an important but undefined B cell contribution to determining GC outcome. As MOG is a tissue-restricted self-Ag, is it unclear how peripheral tolerance mechanisms studied in the context of systemic self-Ag could influence the MOG GC. Indeed, Wang *et.al*. (13) recently showed that MOG-sp. B cells are deleted exclusively in the unique skull bone marrow/leptomeninges developmental niche that has direct access to the CNS where MOG is expressed. However, MOG-sp. B cells were present in apparently normal frequencies in the periphery. Therefore, outside of this niche-specific example of systemic tolerance, as yet there is no indication that these B cells specific for a tissue-restricted Ag undergo peripheral tolerance.

Here, we visualized pre-GC interactions between T and B cells participating in the response to MOG immunization and found that they were of lower quality than those driving the response to the model foreign Ag NPOVA. Further analysis revealed that MOG-sp. B cells failed to upregulate important molecules that mediate interactions with T cells, and that these B cells are inherently less responsive to BCR stimulation, despite the absence of typical indicators of B cell tolerance such as low surface IgM expression.

## Materials and Methods

### Mice

C57Bl/6, 2D2 TCR-transgenic (17), SMARTA TCR-transgenic (4694; Tg(TcrLCMV)327Sdz/JDvsJ), and OTII TCR-transgenic mice (4194; Tg(TcraTcrb)425Cbn/J) were purchased from Jackson Laboratories. B1-8 mice (32) with a homozygous deletion of the Jκ locus (33) were a generous gift from Dr. Ann Haberman. IgH^MOG^ MOG-specific BCR knockin mice (34) were received as a gift from Dr. H Wekerle. Mice expressing fluorescent proteins within all nucleated cells, either dsRed (RFP; 6051; Tg(CAG-DsRedpMST)1Nagy/J) under control of the β-Actin promoter or eGFP via the ubiquitin promoter (4353; Tg(UBCGFP)30Scha/J) were obtained from the Jackson Laboratory. Mice were housed in a specific pathogen-free barrier at West Valley Barrier. Healthy recipient mice between the ages of 6-12 weeks and donors between the ages of 6-16 weeks of age were used in experiments. Mice were sex-matched between groups of an experiment and for mice used in transfers. Cages were split between groups to minimize differences in age. Both male and female mice were used with no apparent differences between the two sexes (data not shown). All of the animal experiments were conducted in compliance with the protocols (2015-090, 2019-123) approved by the Western University Animal Use Subcommittee.

### Antibodies and Reagents

The following Abs were purchased from BD Biosciences: anti-CD4-V450 (RM4-5), anti-CD19-BV711 (1D3), and anti-ICOS-BV421 (7E.17G9) −14). The following Abs were purchased from BioLegend; anti-CD11a-AL488 (M17/4), anti-ICAM1-PECy7 (YN1/1.7.4), anti-SLAM (CD150)-PECy7 (TC15-12F12.2), and anti-Histag-A647 (J099B12). The following Abs were purchased from eBioscience: anti-IgD-eF450 (11–26c), anti-IgD-APC (11–26c), and anti-CD4-PE-Cy5 (RM4-5). Anti-ICOSL-PE (HK5.3) and Fixable Viability Dye eFlour 506 was purchased from Life Technologies. Fluorescently-labeled mMOGtag directly conjugated to Alexa Fluor 647 was generated in house using the NHS Ester fluorophore labeling kit (Life Technologies) according to the manufacturer’s instructions.

### Antigen Reagents

mMOG_tag_ protein was generated as previously described (15, 35). Briefly, the mMOG_tag_ protein is composed of the extracellular domain of mouse myelin oligodendrocyte glycoprotein (MOG1-125) fused to thioredoxin to improve solubility. pET-32a(+) mMOG_tag_ was expressed in BL21 E. *coli* and protein was isolated and purified on His-binding nickel resin (Novagen, Madison, Wisconsin). Nitrophenyl-haptenated ovalbumin (NP-OVA, 25:1 ratio) or NP-MOG (25:1 ratio) were generated in house using NP-OSu (Biosearch Technologies Inc.) and either Ovalbumin protein (Sigma) or mMOG_tag_ protein.

Multivalent MOG-bead antigens were generated using non-functionalized 1 μm diameter silica beads (Bangs Laboratories, Inc. (SS04000) using the manufacturer’s protocol. Briefly, 10 μL of the 10% solids bead solution was washed using 1 mL of acetate buffer (pH 4.6). Following this, beads were resuspended with a concentration of 1 mg/mL or 10 mg/mL of mMOG_tag_ in 100 μL of acetate buffer. Incubations were carried out at room temperature for 90 min with gentle mixing. Beads were washed and resuspended to a final concentration of 6 x 10^9^ beads/mL using the acetate buffer. Beads were stored at 4°C and used within 4 weeks of production.

### Adoptive transfer of cells and immunization

Naïve antigen-specific T cells were isolated from RFP^+^ 2D2 and OTII mice and naïve antigen-specific B cells were isolated from GFP^+^ IgH^MOG^ and B1-8 Jκ^-/-^ mice as previously described (36). Briefly, lymph nodes and spleens of RFP^+^ antigen-specific T cell and GFP^+^ antigen-specific B cell mice were dissociated in ice cold easysep buffer (1mM EDTA, 2% FBS in PBS) and B and T cells were isolated using EasySep Negative selection Mouse B and T cell Enrichment Kits according to the manufacturers protocol (StemCell Technologies). Cells were then suspended in transfer buffer (10 mM HEPES, 25 μg/mL gentamycin, 2.5% Acid citrate dextrose solution A in PBS) and unless otherwise stated, 5 × 10^5^ RFP^+^ T cells and either 1 x10^6^ GFP^+^ B1-8 Jκ^-/-^ or 5 × 10^6^ GFP^+^ IgH^MOG^ B cells (to account for the fact that only 20% are MOG-specific (27)) were transferred i.v into SMARTA recipients 2 d prior to immunization. In intravital experiments using CellTracker Deep Red (ThermoFisher), 5×10^7^ naïve C57Bl/6 B cells were labelled according to the manufacturer’s protocol prior to transfer and imaging. Mice were immunized in the right footpad for intravital experiments or the left and right base of tail with equimolar amounts of the given antigen (125 μg mMOG_tag_, 175 μg NPOVA, 125 μg NPMOG_tag_ (both at a 1:25 protein:NP ratio)) emulsified in CFA at a 1:1 ratio by sonication. Unless otherwise stated, draining popliteal (footpad) or inguinal (base of tail) lymph nodes were harvested at the indicated time points for analysis.

### Multiphoton Intravital Microscopy

At 2 days post-immunization, the popliteal lymph nodes of anesthetized mice were imaged. Mice were initially anesthetized by intraperitoneal injection of a ketamine and xylazine mixture and subsequently with inhaled isoflurane. Animals were immobilized on a custom-built stage with heating mat (Harvard Apparatus) and the right popliteal lymph node was surgically prepared as described previously (37). The lymph node was immersed in saline and covered with a glass coverslip. Temperature of the lymph node was maintained at 37°C during imaging.

Images were acquired on a TRIM II LaVision Biotech (Germany) multiphoton confocal microscope equipped with a Spectra-Physics Insight Dual-Beam laser. For 4D analysis of cell migration, stacks of 19 optical sections with 3 μm z spacing were acquired every 20 s for 60–120 min with the laser tuned to a wavelength of 860 nm. Each xy plane spanned 512 μm in each dimension with a resolution of 0.8 μm per pixel.

### Multiphoton Imaging Analysis

All movies are displayed as 2D maximum intensity projections, unless otherwise stated. Imaris software (Bitplane/Oxford Instruments) was used compile movies and B and T cell interactions were identified in 3D-views and interaction duration determined. Individual B and T cells were tracked using the Imaris cell tracking module using the spot function. All cell tracks were individually examined to confirm that they were complete and reported the behavior of a single cell. The T cell– B cell surface entanglement index was analysed on z-projection images with manual outline of the T cell perimeter and T cell–B cell contact interface, measurements were done using ImageJ.

### Flow cytometry

For analysis of immune responses *in vivo*, draining inguinal lymph nodes were harvested from immunized mice. For analysis of *ex vivo* simulation of B cells, cells were isolated from inguinal, brachial, and axillary lymph nodes using EasySep™ Mouse B Cell Isolation Kit, as described above. 5 × 10^5^ B cells were then incubated for 4 h in RPMI alone (unstim) or with the addition of 15 × 10^5^ low, hi MOG beads, or 20 *μ*g/mL soluble Anti-IgM at 37°C + 5% CO_2_.

Following isolation, cells were prepared for FACS analysis as previously described (15). Briefly, cells were incubated with anti-Fcγ receptor, CD16/32 2.4G2 (BD biosciences), in FACS buffer (2% FBS in PBS) for 30 mins on ice before further incubation with the indicated antibodies. Cells were then stained on ice for 30 mins with the antibodies listed above followed by a secondary stain with streptavidin, or anti-His-A647 for 15 mins in ice, if needed. Dead cells were identified by staining with the Fixable Viability Dye eFluor506 (eBioscience) according to manufacturer’s protocol. Cells were washed with FACS buffer between separate stains. For *ex vivo* experiments, cells were fixed for 20min in 5% PFA. Flow cytometry was performed on a BD Immunocytometry Systems LSRII cytometer and analyzed with FlowJo software (Treestar).

### RNA Isolation and Quantitative PCR

Total RNA was isolated from sorted MOG-sp. and non-MOG-sp. IgH^MOG^ B cells using Trizol^®^ reagent (Life Technologies Inc. Burlington ON) and cDNA was synthesized using Superscript IV VILO Mastermix (Life Tech). Duplex qPCRs were performed using TaqMan Fast Advanced Master Mix (Life Tech), a QuantStudio™ 5 Real-Time PCR System, and the following Taqman Probes (ThermoFisher): *Icam1* (Mm00516023_m1), *Ets1* (Mm01175819_m1), *Nfatc2* (Mm01240677_m1), and *Ywhaz* (Mm03950126_s1). *Icam1, Ets1*, and *Nfatc2* probes were labelled with FAM fluorescent dye and *Ywhaz*, VIC-primer-limited (VIC_PL) assay was used as a normalization control. Relative gene expression was analyzed by the ΔΔCt method.

### Statistical Analysis

PRISM software was used to analyze all FACs and multiphoton data. For statistical comparisons, a Student’s t-test was used for single comparisons and a one-way ANOVA followed by a T test with Bonferroni correction was used for multiple comparisons. No blinding or randomization was used to assign mice to groups. Mice were excluded from experiments if the cell transfer was noted to be unsuccessful at the time of injection. Information about the statistics used for each experiment can be found in the figure legends. Each data point shown represents a single mouse unless otherwise stated.

## Results

### Intravital microscopy of B cell / T cell interactions in immune responses with different B cell outcomes

To directly observe pre-GC interactions between B cells and T cells that naturally generate different outcomes with respect to B cell activation and fate choice, we performed multiphoton confocal intravital microscopy of draining lymph nodes of immunized mice, as previously described (36). RFP-fluorescent T cells (either OVA_329-227_-specific OTII T cells or MOG_35-55_-specific 2D2 T cells) were transferred along with GFP-fluorescent B cells (either NP-specific B18 Jκ^-/-^ or MOG-specific IgH^MOG^ GFP^+^ B cells) to non-fluorescent recipient mice expressing a transgenic TCR to an irrelevant peptide (SMARTA mice). This approach excludes an endogenous T cell response to Ag and ensures that all T cells participating in the response can be identified by RFP-fluorescence. Two days post transfer, recipient mice were immunized in the footpad with the relevant antigen (either NPOVA or MOG_tag_ in CFA). Draining popliteal lymph nodes were imaged 2d post immunization to observe interactions between cognate T and B cells at the B cell Follicle and T cell Zone.

Consistent with previous observations from ourselves and others (36–38), fluorescent Ag-sp. T cells and B cells were apparent in the interfollicular zone at the 2d post immunization timepoint in the NPOVA model system (Figure 1A, Movie 1). This was also the case in the MOG-antigen system (Figure 1A, Movie 2), and there was no significant difference in T cell or B cell migration velocities between model systems (Data Not Shown). Interacting Ag-sp. T and B cells were identified manually and interaction duration was determined by identifying the start of the interaction based on either the first frame of cell:cell contact or first appearance of the interacting cells in the video, and the subsequent end of the interaction based on either the dissolution of the contact or the cells moving out of the field of view (Movies 3 and 4). The average duration of interactions was significantly shorter in the MOG-Ag system compared to the NP-OVA-Ag system (Figure 1B). This difference is almost certainly an underestimate, as it was rare to observe a whole interaction from beginning to end in the NPOVA-Ag system and therefore many interactions lasted much longer than measured. In contrast, because short-duration interactions dominated in the MOG-Ag system, it was much more common to observe an entire interaction between two cells. Indeed, very few MOG-driven interactions lasted longer than 15 min, with most lasting fewer than 5 min (Figure 1C), while only 20% of NP-OVA-driven interactions were this short. Consistent with these observations, the membranes of interacting MOG-sp. T and B cells were less entangled than those of interacting T and B cells in the NPOVA response (Figure 1D), reflecting less engagement of the cells. This has been linked to the strength of signalling between interacting cells (38). Combined, these observations suggest that interactions between MOG-sp. T and B cells are of lower quality, potentially with less exchange of critical activation and differentiation signals, compared to interactions in the standard NPOVA-model system.

**Figure 1:**
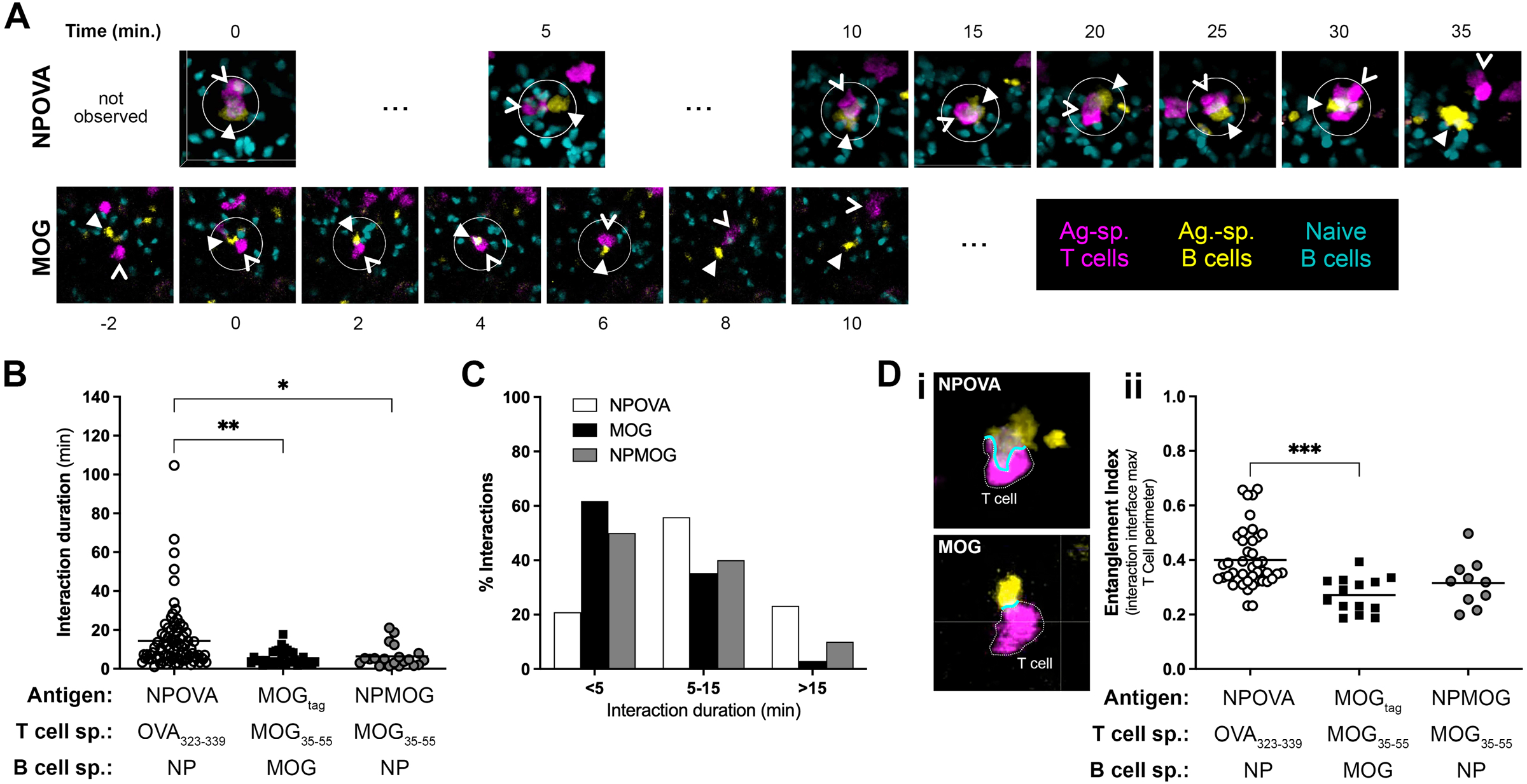
Interactions between MOG-sp. T and B cells are shorter and less entangled that those between T and B cells responding to the model antigen NPOVA. Antigen-specific RFP^+^ T and GFP^+^ B cells were isolated and transferred to SMARTA recipient mice. Recipient mice were immunized 2d later with NPOVA, MOG_tag_, or NPMOG_tag_ in CFA, as indicated. Intravital microscopy of draining popliteal lymph nodes was performed 2d post immunization. (**A**) Still frames from representative 3D videos showing a typical interaction between T (open arrow) and B cells (closed arrow) responding to NPOVA (top) or MOG_tag_ (bottom) immunization. Note that the T/B interaction in the NPOVA system initiated prior to the beginning of the video, as was common for this model. The interaction was observed for a further ~35min before they separated (see Movie 3). The MOG T/B interaction shown is representative of the most common short interactions in this model system lasting less than 10min (see Movie 4). (**B**) Interactions were identified manually and their duration determined by identifying the timepoint when the cells first made contact or came into view, and then when they separated or left the field of view. (**C**) Interaction durations were binned between short (<5 min), medium (5-15 min), and long (>15 min). (**D**) Membrane entanglement was determined by identifying a frame with greatest contact between the interacting T and B cell. (***i***) The length of the membrane contact between the T and B cells was determined and expressed as ratio of the full T cell peripheral measurement (*ii*). Each symbol represents an individual interaction with data pooled from 4 independent experiments for each Ag group. P <0.05, **p <0.01, ***p <0.001, based on an ANOVA followed by a Students *t* test with Bonferroni correction was used for multiple comparisons.

In our previous analysis of the MOG-driven GC response (9), we showed that TCR affinity for Ag was at least partly responsible for the differences in outcome of pre-GC interactions. To determine if the T cell partner controls the quality of the cognate interactions with B cells, we generated a hybrid model system that pairs NP-sp. B cells with MOG-sp. T cells by NP-haptenating MOG_tag_ protein, as previously described (9). Interactions between NP-sp. B1-8 Jκ^-/-^ B cells presenting antigen to MOG-sp. 2D2 T cells resulted in an intermediate interaction duration compared to the long NPOVA-driven and short MOG-driven interactions (Figure 1 B and C), and membrane entanglement was similarly intermediate (Figure 1D). This suggests that the T cell partner is only partly responsible for determining the quality of the T/B interaction. This is consistent with our previous observation that placing NP-sp. B cells under the control of MOG-sp. T cells did not fully recapitulate all aspects of the defective anti-MOG GC response (9).

### Surface molecules that mediate interactions between T and B cells are differentially expressed in the model antigen systems

In separate experiments, fluorescent, Ag-sp. T and B cells were transferred to non-fluorescent SMARTA mice as described above. Recipients were immunized with the appropriate NPOVA, MOG_tag_, or hybrid NPMOG_tag_ antigen and draining lymph nodes were harvested 2d later for analysis by Flow Cytometry. Responding T cells were identified as RFP^+^ CD62L^lo^ while responding B cells were identified as GFP^+^ IgD^lo^. The expression of molecules known to facilitate T/B cell interactions was measured on these cells and normalized to non-fluorescent, endogenous naïve CD62L^hi^ T cells or IgD^hi^ B cells, respectively.

The expression of ICOS, a surface receptor previously shown to be important for communication with B cells (38), was increased 7-8 fold on T cells responding to NPOVA, MOG_tag_, or NPMOG_tag_ compared to naïve cells (Figure 2A*i*). Expression of CD11a, an integrin that contributes to the immune synapse (39), did not change from baseline (Figure 2A*ii*) while the expression of CD150, a molecule previously shown to be important for stabilizing interactions between T and B cells through the adaptor protein SAP (5, 7, 40) increased by 2-3 fold on antigen-responding cells (Figure 2A*iii*). Importantly, we did not observe any differences in T cell expression of these molecules between the three different Model Ag Systems.

**Figure 2:**
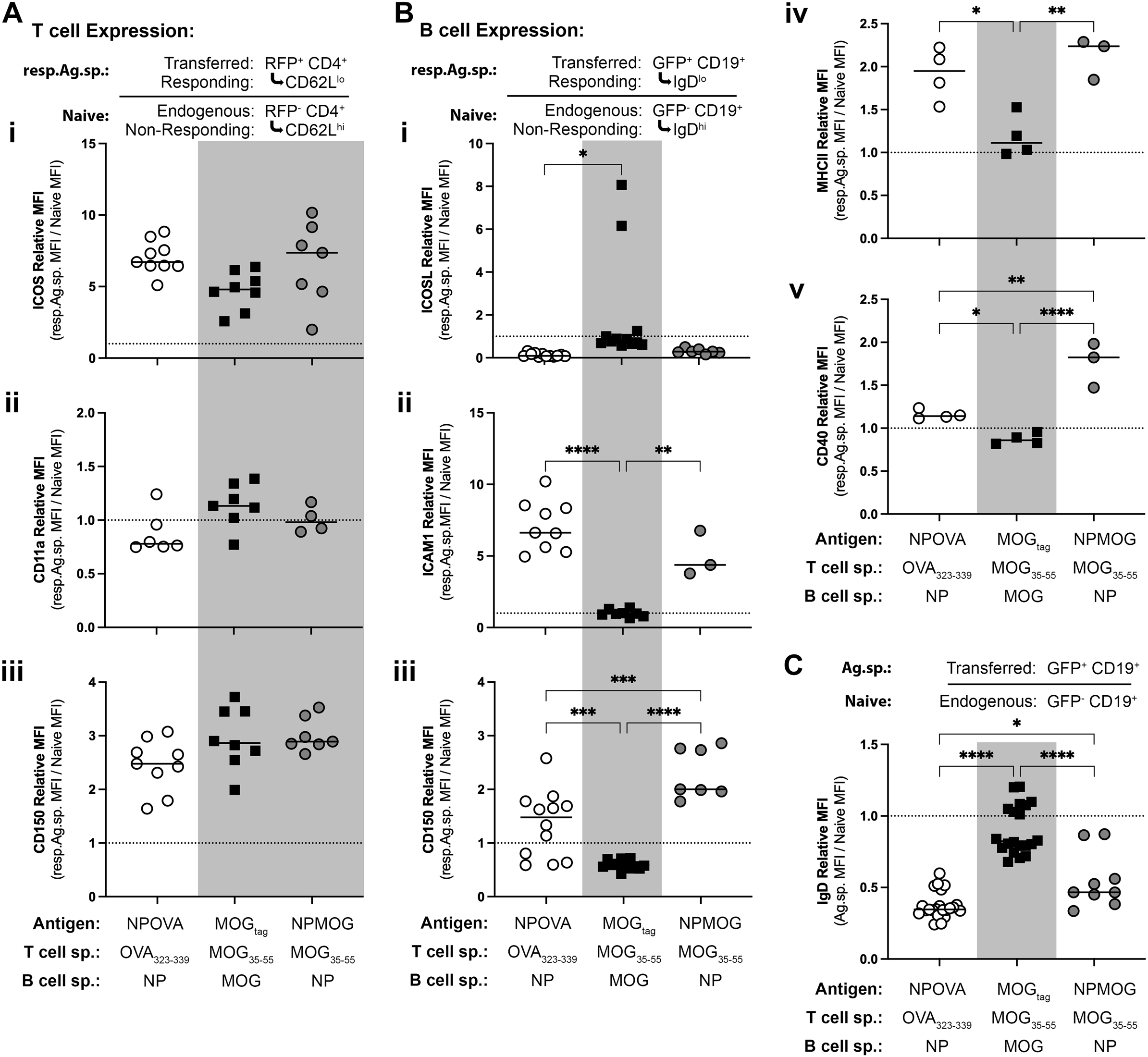
Expression of molecules that mediate T/B interactions on antigen-activated T and B cells *in vivo*. Ag-sp. RFP^+^ T and GFP^+^ B cells were isolated and transferred to SMARTA recipient mice, which were immunized subcutaneously 2d later with NPOVA, MOG_tag_, or NPMOG_tag_ in CFA, as indicated. Cells from draining inguinal lymph nodes were harvested 2d later and analyzed by FACS. (**A**) Responding Ag-sp. T cells were assessed for expression of (***i***) ICOS, (***ii***) CD11a, and (***iii***) CD150. Relative expression based on Geometric Mean Fluorescence Intensity compared to endogenous, naïve T cells from the same mouse is shown, with the value of 1 representing naïve expression levels. (**B**) Responding antigen-specific B cells were assessed for the expression of the corresponding receptors (***i***) ICOSL, (***ii***) ICAM-1, and (***iii***) CD150. B cell expression of (***iv***) MHC class II and (***v***) CD40 was also assessed. Relative expression compared to endogenous, naive B cells from the same mouse is shown, with the value of 1 representing naïve expression levels. (**C**) Relative IgD expression levels by all transferred GFP^+^ B cells compared to endogenous GFP^-^ B cells. Results pooled from at least 2 independent experiments for each marker, except for MHC II and CD40 (B *iv* and *v*). Note that not every experiment stained for each marker, resulting in different numbers of points per plot. Each symbol represents an individual mouse. *p <0.05, **p <0.01, ***p <0.001, **** p <0.0001, based on an ANOVA followed by a Students *t* test with Bonferroni correction was used for multiple comparisons.

In contrast, there were clear differences in the expression of the corresponding ligands on responding B cell partners. ICOSL, the ligand for T cell ICOS (38) was lower on NP-sp. B cells regardless of whether they were paired with MOG- or OVA-sp. T cells (Figure 2B*i*). This may reflect ICOSL shedding, which is necessary for proper regulation of ICOS on Tfh (41). Regardless, no downregulation was observed on B cells responding to MOG-Ag. The expression of both ICAM-1 (Figure 2B*ii*), a CD11a ligand in the immune synapse (42), and CD150 (Figure 2B*iii*), which self-partners CD150 on T cells (40) was increased on responding NP-sp. B cells partnered with either MOG- or OVA-sp. T cells, but not on MOG-sp. B cells. Similar analysis revealed that the expression of both MHC Class II (Figure 2B*iv*) and CD40 (Figure 2B*v*) by MOG-responding B cells was significantly lower on MOG-sp. B cells compared to NP-sp. B cells independent of T cell partner.

Together, these finding suggest that the T cell in an interaction with a B cell does not control the expression of important interaction molecules on its B cell partner. Rather, it suggests that MOG-sp. B cells are less responsive to antigen-stimulation than are NP-sp. B cells. Consistent with this, analysis of IgD expression on all transferred GFP^+^ B cells showed that IgD downregulation, a marker of activation, was significantly greater on NP-sp. B cells, regardless of whether partnered with OVA- or MOG-sp. T cells, compared with MOG-sp. B cells (Figure 2C).

### Multivalent antigen does not correct the anti-MOG B cell response

Differences in B cell responsiveness to NP-*vs* MOG Ag could either be due to a difference in the B cells themselves, or because of some difference in the nature of the Ag. NP-haptenation of a protein carrier, in this case either OVA or MOG_tag_, results in a multivalent antigen, which has been previously shown to produce a stronger BCR signal and greater B cell activation (43) compared to a monovalent antigen like soluble MOG_tag_. To determine if epitope valency is responsible for the differences in B cell responsiveness, we generated a multi-valent MOG-Ag by adsorbing different amounts of mMOG_tag_ to 1μm silica beads. Flow cytometry analysis demonstrated the successful generation of beads coated with different levels of MOG_tag_ Ag (Figure 3A). IgH^MOG^ B cells were isolated as described above and cultured for 4hr in the presence of Low or High valency MOG-beads and B cell activation was subsequently assessed by flow cytometry.

**Figure 3:**
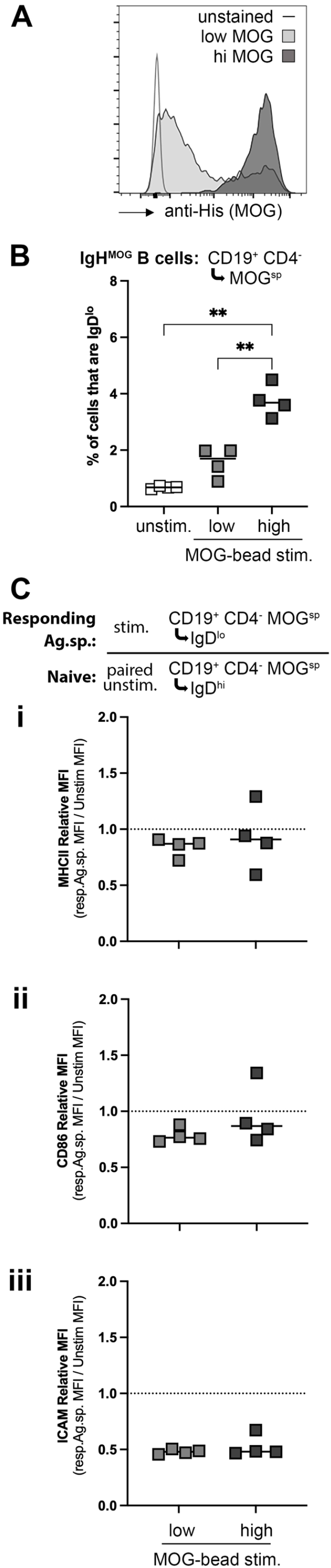
Antigen valency does not correct the poor responsiveness of MOG-sp. B cells. Multi-valent MOG antigen was generated by adsorbing MOG_tag_ protein to 1 μm silica beads. (**A**) anti-His_6_ Tag antibodies were used to measure the amount of MOG_tag_ on low- and high-valency beads (incubated with 1mg/ml or 10mg/ml MOG_tag_, respectively). (**B**) B cells were isolated from IgH^MOG^ mice and incubated for 4 hrs in RPMI alone (unstim) or with the addition of 1.5 x 10^6^ low or hi MOG beads (1:3 cells:beads ratio) at 37°C + 5% CO_2_. The downregulation of IgD on MOG-sp. B cells as a marker of activation was assessed by FACS. (**C**) The relative expression of (***i***) MHC Class II, (***ii***) CD86, and (***iii***) ICAM-1 on responding IgD^lo^ MOG-sp. B cells compared to paired unstimulated MOG-non-sp. B cells is shown. Results from one representative of two independent experiments are shown. Each symbol represents data from cells isolated from an individual mouse. **p <0.01 based on an ANOVA followed by a Students *t* test with Bonferroni correction for multiple comparisons.

Not all B cells with the IgH^MOG^ heavy chain knockin are MOG-sp., as specificity also depends on the light chain. Using a fluorescently labelled MOG_tag_ protein as a reagent to identify MOG-binding B cells by Flow Cytometry, we typically find that ~20% of IgH^MOG^ B cells are MOG-sp. (27). Using this approach to observe the response of MOG-sp. B cells only, we observed a degree of IgD downregulation in response to both High and Low Valency MOG_tag_-bead antigens (Figure 3B), with the High-Valency beads inducing a greater percentage of IgD^lo^ B cells, as anticipated. However, neither the Low- or High-Valency beads induced an upregulation in MHC class II, CD86, or ICAM1 relative to naïve IgD^hi^ MOG-negative B cells (Figure 3C*i ii iii*). This was consistent with the lack of change in these molecules observed *in vivo* on adoptively-transferred B cells responding to MOG immunization (Figure 2). Therefore, increasing the valency of the MOG-antigen does not correct the apparent defect in the responsiveness of MOG-sp. B cells.

### MOG-specific B cells are less responsive to BCR stimulation in vitro

To determine if MOG-sp. B cells are inherently less responsive to BCR stimulation, B cells were isolated from wild-type or IgH^MOG^ mice and incubated with anti-IgM antibodies as an Ag-agnostic BCR stimulation. As expected, anti-IgM induced IgD downregulation in a significant proportion of wild-type B cells compared to unstimulated cells (Figure 4A*i*). In contrast, the proportion of IgD^lo^ IgH^MOG^ B cells in response to anti-IgM was not significantly different compared to unstimulated cells (Figure 4B*i*), suggesting that B cells from IgH^MOG^ mice may indeed be inherently less responsive to BCR stimulation than are wild-type B cells. Using fluorescent MOG_tag_ protein to separately analyze MOG-sp. and non-sp. IgH^MOG^ B cells, we found that a significant number of IgH^MOG^ MOG non-sp. B cells did in fact respond to anti-IgM by downregulating IgD (Figure 4B*ii*), but this was less evident in the MOG-sp. pool (Figure 4B*iii*). Interestingly, a direct comparison of IgD^lo^ B cells showed that anti-IgM-responding MOG-sp. IgH^MOG^ cells expressed significantly less MHC class II and CD86 compared to responding non-MOG-sp. cells (Figure 4 D*i ii*).

**Figure 4:**
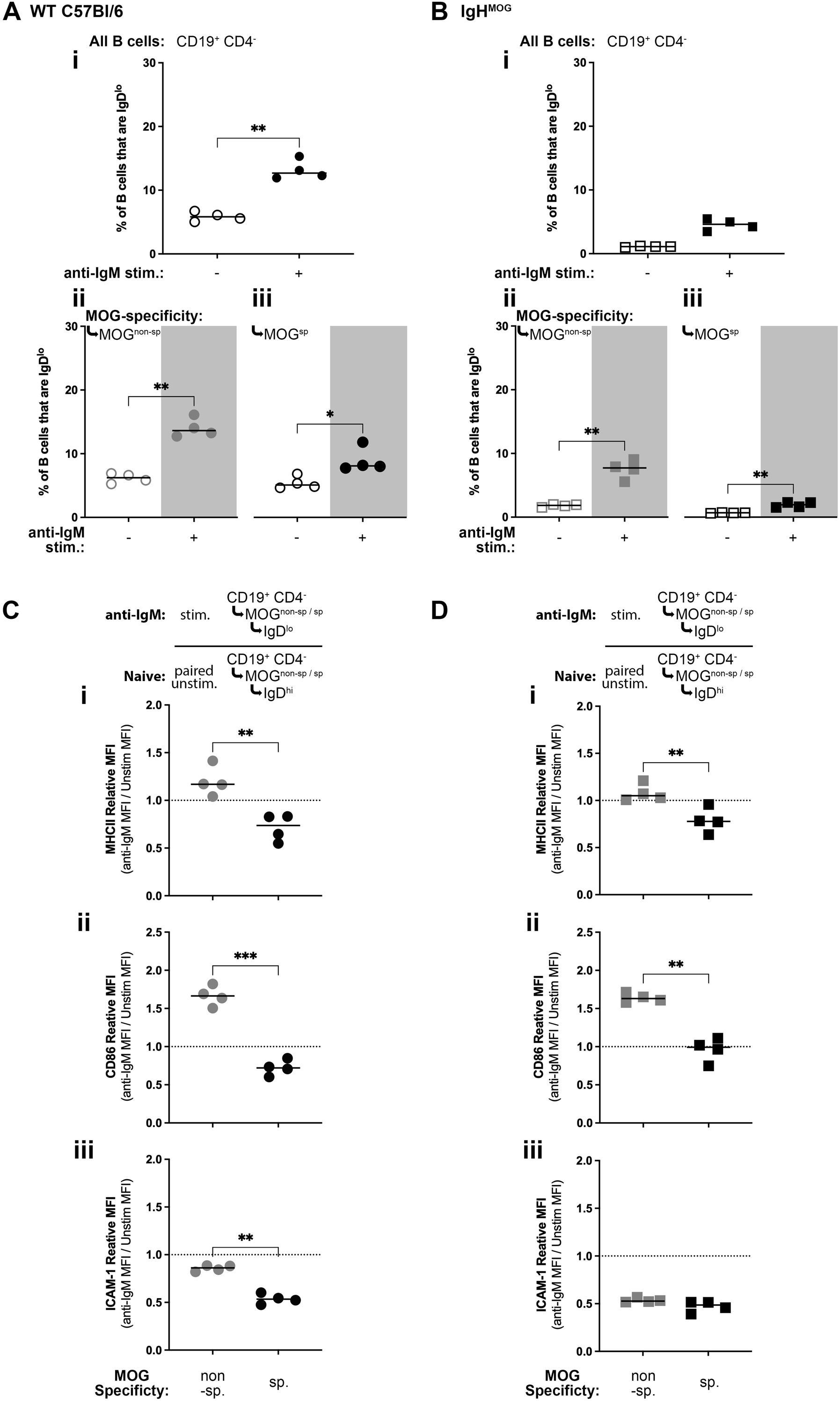
MOG-sp. B cells are inherently less responsive to BCR stimulation. B cells were isolated from wild-type C57Bl/6 or IgH^MOG^ mice and incubated for 4 hrs in RPMI alone (unstim) or with 20 μg/ml anti-IgM antibodies. The downregulation of IgD on (**A**) wild-type or (**B**) IgH^MOG^ B cells was assessed by FACS for (***i***) all CD19^+^ B cells, or separated into (***ii***) non-MOG-sp. and (***iii***) MOG-sp. B cells. Responding Non-MOG-sp. and MOG-sp. (**C**) Wild-Type and (**D**) IgH^MOG^ B cells were further assessed for the expression of antigen-presentation-associated surface molecules (***i***) MHC Class II, (***ii***) CD86, and (***iii***) ICAM-1. Expression levels are shown as relative to paired non-MOG-sp. or MOG-sp. unstimultated, naive B cells, as appropriate. One representative of two independent experiments is shown. Each symbol represents data from cells isolated from an individual mouse. *p <0.05, **p <0.01, ***p <0.001 based on an ANOVA followed by a Students *t* test with Bonferroni correction for multiple comparisons.

Using the same fluorescent MOG_tag_ labeling approach, we similarly compared the much rarer MOG-sp. B cells from wild-type mice to the majority MOG-non-sp. B cells following anti-IgM stimulation. As for MOG-sp. IgH^MOG^ B cells (above), responding IgD^lo^ MOG-sp. wild-type B cells also expressed significantly less MHC class II and CD86 on their surface compared to IgD^lo^ MOG-non-sp. B cells (Figure 4C*i ii*). On wild-type MOG-sp. B cells, ICAM-1 expression was significantly lower compared to non-sp. B cells (Figure 4C*iii*), while this phenotype was inconsistent on IgH^MOG^ cells, in that there was no significant difference in the experiment shown here (Figure 4D*iii*), but ICAM-1 expression was significantly lower in a separate independent experiment (Not Shown). Together, this suggests that MOG-specificity itself results in reduced responsiveness to BCR stimulation, and that this is not an artifact of the IgH^MOG^ knockin mutation.

### MOG-specific B cells do not express markers of B cell tolerance

B cell tolerance is associated with reduced surface IgM expression (44–47). We therefore compared IgM levels on unstimulated MOG-sp. and non-sp. IgH^MOG^ B cells. No decrease in surface IgM was noted on MOG-sp. B cells and, in fact, IgM levels were higher on MOG-binding cells (Figure 5A). Both Ets-1 (48) and NFAT1 (49) are transcription factors that have been shown to be necessary for B cell anergy induced by repeated exposure to systemic autoantigen. We therefore sorted MOG-sp. and non-sp. IgH^MOG^ B cells from lymph nodes and isolated mRNA for analysis of the expression of these genes by qPCR. Both MOG-sp. and non-sp. B cells expressed these transcription factors at the same level (Figure 5B*i ii*). Upregulated ICAM-1 expression on B cells has been shown to be associated with and necessary for anergy induced in mice treated with anti-CD45RB to prolong allograft survival (50). While we did not observe any evidence for transcriptional upregulation of ICAM-1 on MOG-sp. B cells (Figure 5B*iii*), flow cytometry of unstimulated B cells freshly harvested from IgH^MOG^ mice did reveal that MOG-sp. B cells expressed significantly more ICAM-1 on their surface compared to non-MOG-sp. B cells (Figure 5C). Therefore, the phenotype of less-responsive MOG-sp. B cells do not have a phenotype that is consistent with most previous descriptions of B Cells anergised via exposure to Ag, but may be more related to anergy induced via therapeutic intervention.

**Figure 5:**
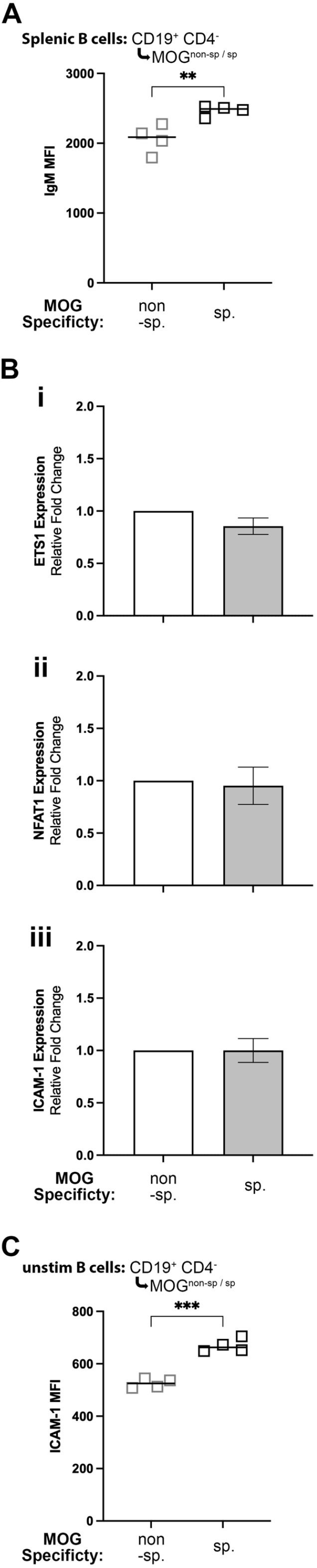
MOG-sp. B cells do not have a phenotype previously associated with B cell tolerance. (**A**) Splenic B cells from naïve IgH^MOG^ mice were assessed for IgM expression by FACS. Geometric Mean Fluorescence Intensity is shown. Each symbol represents data from cells isolated from an individual mouse. **p <0.01 based on a paired Students t test. (**B**) Non-MOG-sp. and MOG-sp. B cells were sorted from IgH^MOG^ mice and mRNA as isolated for analysis of (***i***) ETS1, (***ii***) NFAT1, and (***iii***) ICAM-1 by qPCR. Expression is shown normalized to non-MOG-sp. cells. (**C**) FACS measurement of surface ICAM-1 expression on non-MOG-sp. and MOG-sp. IgH^MOG^ B cells was assessed by FACS. Geometric Mean Fluorescence Intensity is shown. Each symbol represents data from cells isolated from an individual mouse. ***p <0.001 based on a paired Students *t* test.

## Discussion

Interactions between cognate T and B cells are integral to the GC response because the signals exchanged during the interactions determine survival and influence the fate of individual B cell clones. Because of this, the nature of these interactions influences the size and quality of the humoral response as a whole to any given Ag. Indeed, studies manipulating molecules known to facilitate these interactions support a model where the quality of the T/B interaction is directly related to the outcome. For example, deletion of the signaling molecules SAP, which have been described to stabilize T cell interactions with B cells but not with other Ag presenting cells (7), resulted in a failed GC response, while deletion of CD150 or other family members that signal through SAP had intermediate outcomes (51). Similarly, the degree of membrane engagement between two interacting T and B cells, presumably a poxy measure of the degree of signal exchange, has been linked to the quality of the response (38). Here, we used two model Ag systems that naturally generate different GC outcomes without external interference to determine if outcome is linked to the quality of the interactions between cognate T and B cells. Our findings suggests that these interactions are indeed a natural checkpoint used by the immune system to control the quality of the GC response. The quality of these interactions is likely to be determined at least in part by BCR-driven alterations in the expression of molecules that facilitate B cell presentation of Ag to, and interactions with, activated T cells.

Our previous analysis of GC outcomes (9) used the same Model Ag Systems and focussed primarily on the outcomes of interactions that occur at later timepoints within the mature GC. At this stage, interactions with Tfh control GC B cell survival and cyclic fate choices between further proliferation or differentiation into either Memory B cells or effector Plasma Cells. In this study, we found that the different outcome of these interactions between Ag Systems were at least partly attributed to T cell-associated factors. For example, the anti-NP GC was smaller under the control of MOG-sp. T cells compared to when driven by OVA-sp. T cells, largely due to preferential B cell differentiation to memory-phenotype or Plasma Cells at the expense of returning to the GC dark zone for additional rounds of proliferation. Further, the size of the anti-MOG GC response could be partly restored by manipulating the antigen to increase TCR affinity. However, not all defects in the anti-MOG GC could be attributed to the T cell partner, and this was particularly true of fate outcomes driven by early pre-GC interactions.

In the current study, we extended the above findings by directly observing the critical pre-GC T/B interactions in these Model Antigen Systems. Our observations are consistent with the hypothesis that the quality of these interactions influences B cell fate choice. Further, because these differences occurred in the absence of external manipulation, they are consistent with the hypothesis that these interactions are natural immune checkpoint to determine the quality of the response to a given antigen. Differences between Model Ag Systems correlated with differences in B cell expression of molecules that facilitate interactions with T cells. Following 2d of Ag stimulation *in vivo*, NP-sp. B cells upregulated surface CD150, along with ICAM-1 and MHC class II compared to naïve cells. There was little evidence that T cells modulated the expression of the associated ligands in response to different Ags, either at the early 2d timepoint investigated here or at the mature GC timepoint, as we reported previously (9). In contrast, the expression of these molecules remained largely unchanged on MOG-sp. B cells. Similarly, *ex vivo* stimulation of either wild-type or IgH^MOG^ B cells revealed that MOG-sp B cells respond differently to short-term (4hr) stimulation in response to Ag-agnostic anti-IgM BCR stimulation, again reflected by lower expression of interaction molecules on MOG-sp. B cells. These suggest that changes in interaction molecule expression are triggered by BCR stimulation, rather than because of instructions received from interacting T cells. It also demonstrates that differences in these signals have consequences to the outcome of the humoral response, and that MOG-sp. B cells themselves are inherently less responsive than NP-sp. B cells or the general naïve B cell population as a whole.

While MOG is a self-Ag and mechanisms of autoimmune B cell tolerance have been described (10, 11), we originally suspected that the difference in B cell responsiveness between MOG- and NP-sp. GCs was driven by properties of the Ags themselves and not, as we now suspect, anergy of the MOG-sp. B cells. We require MOG-deficient mice to demonstrate this formally, but we have not been able to acquire them. The reason that we did not originally think that MOG-sp. B cells are tolerized is that, using a strain of MOG-deficient mice, Delarasse *et.al*. (52) did not observe any evidence of differences in T or B cell immune responses to MOG immunization compared to wild-type mice. In contrast, using a different MOG-deficient strain Linares *et.al*. (53) observed evidence of T cell tolerance to MOG but did not test the B cell response. More recently Wang *et.al*. (13) performed an elegant series of experiments in non-human primates and mice, including IgH^MOG^ mice, that clearly demonstrate central tolerance mechanisms deleting MOG-sp. B cells exclusively from the unique leptomeninges B cell developmental niche that has direct access to the CNS where MOG is expressed. However, as we show here and has been demonstrated previously (27, 34), they also found that MOG-sp. B cells remain highly enriched in the peripheral lymphatics of IgH^MOG^ mice, suggesting that tolerance mechanism were limited to sites where B cells are exposed to self-Ag during development. Mechanisms of B cell tolerance and especially peripheral tolerance are generally considered to be less stringent compared to that for T cells, in part because many responses depend on T cell help and therefore benefit from T cell regulatory mechanisms, and also because B cell Ag recognition is not MHC restricted and not constrained to a narrow range of affinities.

Our strongest evidence that MOG-sp. B cells in the periphery have undergone some tolerance process is the differential responsiveness of MOG-sp. *vs*. MOG-non-sp. B cells to anti-IgM stimulation, whether they were isolated from wild-type or mutant IgH^MOG^ mice. Further, increasing the valency of the MOG Ag failed to rescue responsiveness, as would be expected if properties of the Ag were responsible for the observed differences. However, this differential response was not due to downregulation of surface IgM which was, if anything, higher on MOG-sp. cells. IgM downregulation has been identified as an important peripheral B cell tolerance mechanism in models of systemic self-Ags (46, 47, 54). B cell exclusion from the follicle has also been described in these systems (55), but we readily observed transferred MOG-sp. B cells in follicles in our intravital imaging studies and extensive previous histology studies (9). Nevertheless, a recent study by Brooks *et.al*. (56) observed that, in a transgenic mouse system where an otherwise-foreign Ag (hen egg lysozyme (HEL)) was expressed ubiquitously, the anti-HEL GC bore some resemblance to the anti-MOG GC in that it was not sustained, and had limited Memory B cell and Plasma Cell differentiation. Therefore, while MOG-sp. B cells do not phenotypically resemble the IgM^lo^ B cells associated with systemic self-Ag, there may be an underlying mechanism limiting the response. We did find the ICAM-1 protein (but not transcription) was upregulated on the surface of naïve MOG-sp. B cells. This was counterinitiative, as this molecule is upregulated on BCR-stimulated cells to facilitate interactions with T cells. However, the ICAM-1^hi^ phenotype has been described on B cells in mice treated with anti-CD45RB to induce tolerance and promote allograft survival (57) and ICAM-1 was shown to be necessary for reduced B cell responsiveness to the allograft in these mice (50).

This may therefore represent a different pathway to limit B cell responsiveness that is independent of regular exposure to Ag. As the B cell response was not corrected by increasing Ag avidity, nor by bypassing BCR affinity for Ag by stimulating with anti-IgM, features of the Ag itself are not responsible for the different outcomes. As surface levels of BCR on MOG-sp. B cells is not reduced, this is not the mechanism to limit B cell activation. Instead, these cells do not respond to BCR-stimulation by altering their expression of molecules to facilitate interactions with cognate T cells at the follicular border. Compared to interactions between T and B cells these interactions are of lower quality and, while they are apparently able to support a reduced degree of B cell clonal expansion, they are not able to support early plasmablast differentiation or class switch.

## Acknowledgements

The authors would like to thank the veterinarians and animal care staff at the West Valley Barrier Facility for their excellent husbandry of our experimental animals. We are also grateful for the efforts of our Work-Study Undergraduate Students who support our mouse breeding program. This study was funded by a Discovery Grant from the National Sciences and Engineering Council of Canada (NSERC) and a Discovery Research Operating Grant from the the Multiple Sclerosis Society of Canada (MSSOC). KAP is the recipient of an endMS Post-Doctoral Fellowship from the MSSOC. ST was a recipient of an NSERC Undergraduate Summer Research Award.

## Notes

### Competing Interest Statement

The authors have declared no competing interest.

